# Sulfane sulfur post-translationally modifies the global regulator AdpA to influence actinorhodin production and morphological differentiation of *Streptomyces coelicolor*

**DOI:** 10.1101/2022.01.06.475307

**Authors:** Ting Lu, Xiaohua Wu, Qun Cao, Yongzhen Xia, Luying Xun, Huaiwei Liu

## Abstract

The transcription factor AdpA is a key regulator controlling both secondary metabolism and morphological differentiation in *Streptomyces*. Due to its critical functions, its expression undergoes multi-level regulations at transcriptional, post-transcriptional, and translational levels, yet no post-translational regulation has been reported. Sulfane sulfur, such as organic polysulfide (RS_n_H, n≥2), is common inside microorganisms, but its physiological functions are largely unknown. Herein, we discovered that sulfane sulfur post-translationally modifies AdpA in *S. coelicolor* via specifically reacting with Cys^62^ of AdpA to form a persulfide (Cys^62^-SSH). This modification decreases the affinity of AdpA to its self-promoter *P*_*adpA*_, allowing increased expression of *adpA*, further promoting the expression of its target genes *actII-4* and *wblA*. ActII-4 activates actinorhodin biosynthesis and WblA regulates morphological development. Bioinformatics analyses indicated that AdpA-Cys^62^ is highly conserved in *Streptomyces*, suggesting the prevalence of such modification in this genus. Thus, our study unveils a new type of regulation on the AdpA activity and sheds a light on how sulfane sulfur stimulates the production of antibiotics in *Streptomyces*.

**IMPORTANCE:** *Streptomyces* produce myriad of polyketide compounds having (potential) clinical applications. While the database of polyketide gene clusters are quickly expanding, the regulation mechanisms of them are rarely known. Sulfane sulfur species are commonly present in microorganisms with unclear functions. Herein, we discovered that sulfane sulfur increases actinorhodin (ACT) production in *S. coelicolor*. The underlying mechanism is sulfane sulfur specifically reacts with AdpA, a global transcription factor controlling both ACT gene cluster and morphological differentiation related genes, to form sulfhydrated AdpA. This modification changes dynamics of AdpA-controlled gene network and leads to high expression of ACT biosynthetic genes. Given the wide prevalence of AdpA and sulfane sulfur in *Streptomyces*, this mechanism may represent a common regulating pattern of polyketide gene clusters. Thus, this finding provides a new strategy for mining and activating valuable polyketide gene clusters.

## INTRODUCTION

*Streptomyces* spp. are gram positive bacteria with a filamentous form, which colonize a wide range of terrestrial and aquatic niches. The most famous characteristic of *Streptomyces* is the ability to produce myriad of secondary metabolites including antibiotics, antifungal, antiviral, anthelmintic agents, antitumoral drugs, anti-hypertensives, herbicides, and interesting pigments (1-3). Numerous efforts have been spent on searching, identifying, and modifying gene clusters responsible for biosynthesis of these secondary metabolites (4). In contrast, much less energy has been invested in illustrating the transcriptional/translational regulation of these gene clusters. One reason is *Streptomyces* have a complex life cycle that includes sporulation, a vegetative or substrate state, and aerial mycelial growth. The biosynthesis of secondary metabolites is closely linked to the stages of life cycle (5, 6), which makes relative studies challengeable.

AdpA is a transcriptional regulator universally present in *Streptomyces* (7). It locates in the second layer of the A-factor-dependent transcriptional network in *Streptomyces griseus*, the first layer is the A-factor receptor, which activates AdpA expression at the present of A-factor (γ-butyrolactone, a quorum sensing hormone). Therefore, AdpA expression is indirectly controlled by the quorum sensing signal. Aside from A-factor, there are at least three players in AdpA expression regulation, including the master developmental regulator BldD regulating at transcriptional level (8, 9), a *cis*-antisense RNA regulating at post-transcriptional level (10), and a rare tRNA (tRNA^UUA^_Leu_) encoding gene *bldA* regulating at translational level (11). It was also reported that AdpA can be transcriptionally self-inhibited (12). Why regulation of AdpA expression is such multiple and complicated might be owing to that AdpA is a key regulator of both secondary metabolism and morphological differentiation (13). Considering the critical functions it conducts, whether there are other players regulating at different levels of AdpA expression or activity is unclear but worthy of further investigation.

Sulfane sulfur containing compounds, such as persulfide (HSSH and RSSH) and polysulfide (HSS_n_H, S_n_, RSS_n_H, RSS_n_R, n≥2), are commonly present in both eukaryotic and prokaryotic cells (14). In the latest two decades, intensive studies of sulfane sulfur have been performed with mammalian cells because it is found that sulfane sulfur is involved in the regulation of diverse physiological and pathological processes including apoptosis, carcinogenesis, and redox maintenance (15-18). On the other hand, studies of microorganism sulfane sulfur are traditionally focused on its metabolism and its role in the global sulfur cycle (19, 20). In recent years, the physiological functions of sulfane sulfur in microorganisms also attract attentions. For instance, Peng et al. (21) found that sulfane sulfur regulates the expression of virulence factors in *Staphylococcus aureus*, Liu et al (22) reported that sulfane sulfur is involved in photosynthesis regulation in *Synechococcus*. Albeit being noticed, the functions of sulfane sulfur in microorganisms are largely unveiled.

In a previous study, we discovered that sulfane sulfur functions as a signal to activate actinorhodin (ACT) production in *S. coelicolor* M145, a model strain of *Streptomyces*. In addition, the spore formation process is accelerated by endogenously accumulated sulfane sulfur (23). These phenomena suggest that sulfane sulfur affects both secondary metabolism and cell cycle in *S. coelicolor* M145. Based on these findings, we herein studied the underlying mechanism of how sulfane sulfur perform such functions. We found that AdpA is the key media of sulfane sulfur signaling. AdpA senses the level of intracellular sulfane sulfur and adjusts ACT production and spore formation. Even the expression of AdpA itself is affected by sulfane sulfur, i.e. sulfane sulfur is a new regulator of AdpA. Thus, this study unveils one way of how sulfane sulfur signals in *Streptomyces*.

## RESULTS

### AdpA is a key regulator of ACT production and morphological development in *S. coelicolor*

Previous studies reported that AdpA involves in the regulation of ACT production and morphological development in *S. coelicolor* (24, 25). Herein, we constructed an *adpA-*disrupted *S. coelicolor* M145 strain (Δ*adpA*). It exhibited a phenotype of no ACT but high undecylprodigiosin (RED) production when cultured on YBP agar medium (Fig. 1A). Complementary expression of *adpA* gene using a plasmid pMS82*-adpA* (Δ*adpA::adpA*) restored ACT production, while the control, Δ*adpA* harboring an empty plasmid (Δ*adpA::*pMS82), showed no change. In addition, we noticed that both Δ*adpA* and the control showed a bald and non-spore form on YBP medium (Fig. 1A) while Δ*adpA::adpA* restored the spore formation. These results verified that AdpA controls ACT production and morphological development in *S. coelicolor* M145 strain.

**Fig. 1.**
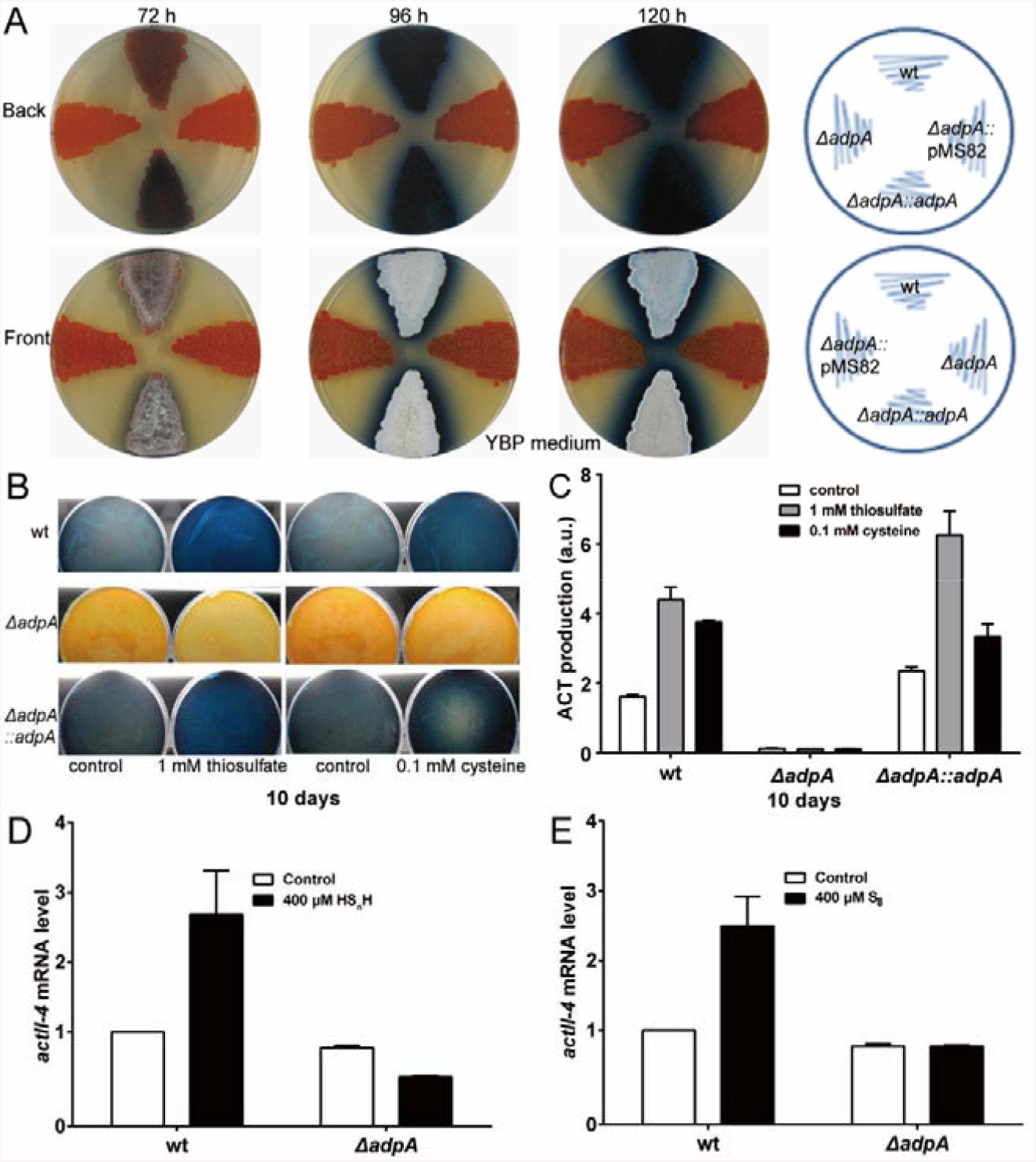
AdpA is required for ACT production and morphological development in *S. coelicolor* M145. (A) Phenotypes of the wt, Δ*adpA*, Δ*adpA::adpA*, and Δ*adpA::*pMS82 strains grown on YBP medium at 30°C. Images were taken at the indicated times. (B) 1 mM thiosulfate or 0.1 mM cysteine were added into YBP agar plates before inoculation. The plates were incubated at 30°C for 10 days, images were captured from the reverse side of the plates. (C) Quantitative determination of ACT produced by the wt, Δ*adpA* and Δ*adpA::adpA* strains on YBP containing thiosulfate or cysteine, the plates were incubated at 30°C for 10 days and then subjected to analysis. Data were from three independent repeats. (D-E) wt and Δ*adpA* strains were grown on YBP liquid medium, when cultured for 36 h, 400 μM HS_n_H or S_8_ were added. After 1 h, RNA samples were isolated after induction.The y-axis shows the fold change of *actII-4* expression levels in wt and Δ*adpA*. Real-time PCR data were from three independent repeats and shown as average ± SD.

### Sulfane sulfur performing ACT activation effect requires the presence of AdpA

Since the Δ*adpA* strain displayed opposite phenotypes as that of sulfane sulfur treated strain (11), we suspected that AdpA had interwound functions with sulfane sulfur. We performed sulfane sulfur induction experiments using the *S. coelicolor* M145 (wt), Δ*adpA* and Δ*adpA::adpA* strains. The strains were spread on YBP medium containing 1 mM thiosulfate or 0.1 mM cysteine (which can be converted to sulfane sulfur *in vivo*) and cultured at 30°C for 10 days. For wt, the production of ACT was significantly increased by thiosulfate/cysteine treatment (Fig. 1B and 1C). For Δ*adpA*, no production of ACT was observed with or without thiosulfate/cysteine treatment. For Δ*adpA::adpA*, the induction effects were similar as in wt (Fig. 1B and 1C). These results indicated that AdpA is required for sulfane sulfur to execute the ACT production activating function.

ActII-4 is the ACT production “pathway-specific” activator (25, 26). We analyzed transcription of *actII-4* using the real-time quantitative reverse transcription PCR (RT-qPCR) method. The wt and Δ*adpA* strains were treated with two sulfane sulfur containing chemicals, hydrogen polysulfides (HS_n_H, n≥2) and sublimed sulfur (S_8_). For wt, the transcription level of *actII-4* was much higher in treated strain than that in untreated one (Fig. 1D and 1E); whereas for Δ*adpA*, the transcription level of *actII-4* had no obvious change after sulfane sulfur treatment (Fig. 1D and 1E). These results indicated that sulfane sulfur can increase ActII-4 expression, which subsequently activates ACT production, but this process requires the presence of AdpA.

### Sulfane sulfur affects the interaction between AdpA and its cognate promoters

AdpA controls the transcription of *actII-4* and *wblA* (*whiB* like gene A, which controls morphological development in *S. coelicolor*) via binding to their promoters (27). Using these two promoters and an enhanced-green-fluorescence-protein encoding gene (*egfp*), we constructed two reporter systems (Fig. 2A and 2B). These reporter systems were introduced into the wt and Δ*adpA* strains. HS_n_H was used to treat the strains containing the reporter systems. After 30 min treatment, the mycelium was collected by centrifugation and the fluorescence was read by a fluorophotometer. In the wt strain, HS_n_H treatment enhanced the strength of both *actII-4* promoter (*P*_*actII-*4_) and *wblA* promoter (*P*_*wblA*_), evidenced by the increased EGFP expression. Whereas, in the Δ*adpA* strain, EGFP expression was not increased but decreased after HS_n_H treatment, indicating that HS_n_H treatment lost the enhancing effect on these promoters. These results suggested that sulfane sulfur may affect the interaction between AdpA and its cognate promoters.

**Fig. 2.**
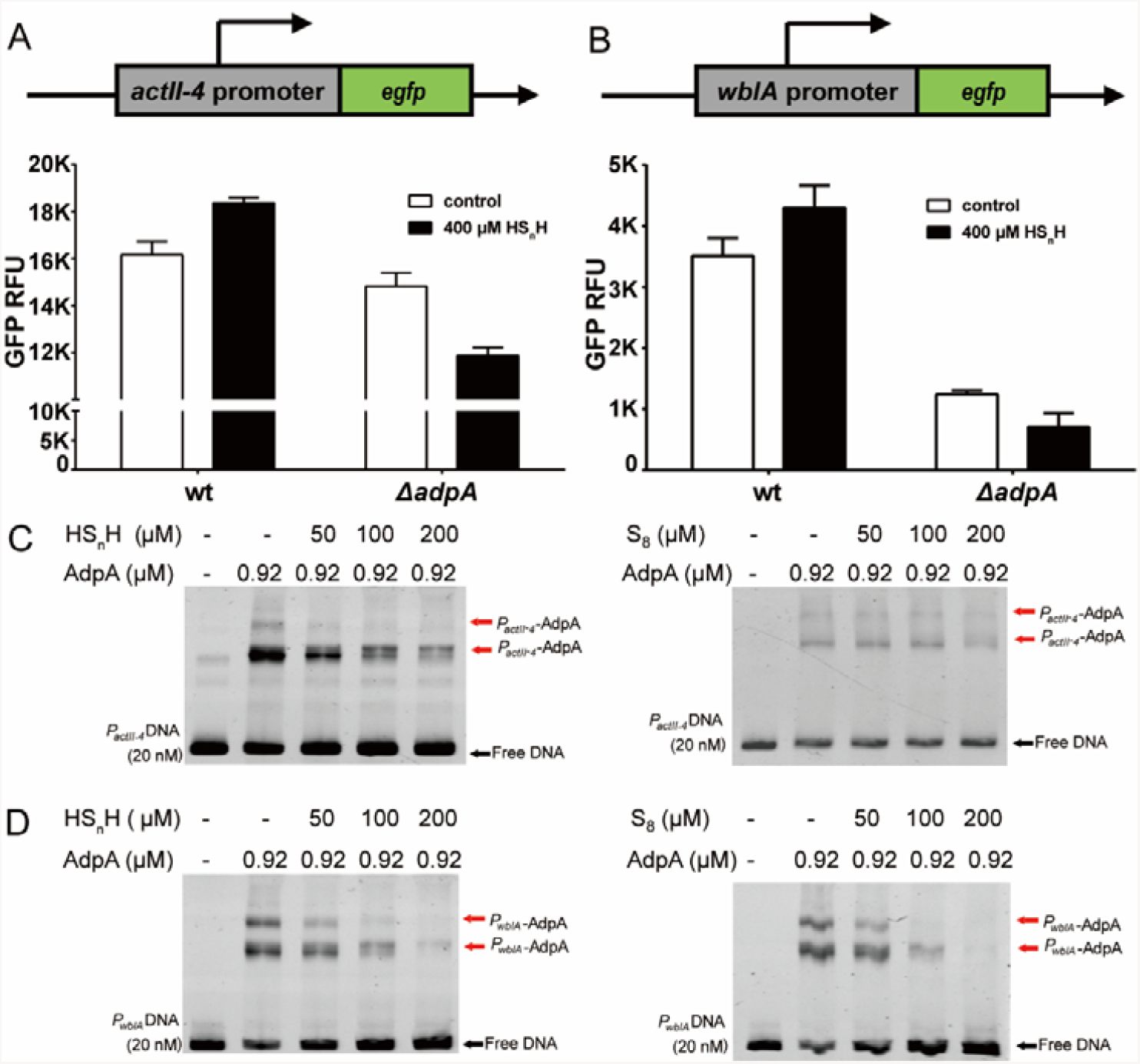
Sulfane sulfur is involved in the process of AdpA regulating target genes. (A) HS_n_H was used to treat wt and Δ*adpA* strains harboring pMS82-*actII-4*p-*egfp*. (B) HS_n_H was used to treat wt and Δ*adpA* strains harboring pMS82-*wblA*p-*egfp*. Data were from three independent repeats and shown as average ± SD. (C,D) EMSA analysis of the AdpA affinity to *P*_*actII-4*_ promoter DNA (C) and the *P*_*wblA*_ promoter DNA (D). All lanes contained 20 nM probe DNA, lanes 2-5 contained the indicated protein concentration, lanes 3-5 contained the HS_n_H (left) or S_8_ (right). Black arrow indicates the free DNA probe and red arrow indicates *P*_*actII-4*_-AdpA or *P*_*wblA*_-AdpA complex.

We then performed electrophoretic mobility shift assays (EMSA) to investigate the interaction. The AdpA protein was expressed in *E. coli* BL21 (DE3) and purified. The DNA probes of the *wblA* and *actII-4* promoters were obtained by PCR. When AdpA was mixed with the *P*_*actII-4*_ or *P*_*wblA*_ DNA probe, it bound to them (Fig. 2C and 2D). When HS_n_H or S_8_ was also added, the fraction of the AdpA-probe complexes decreased (Fig. 2C and 2D). These results indicated that sulfane sulfur decreased the affinity of AdpA to *P*_*actII-4*_ and *P*_*wblA*_.

To test whether the influence of sulfane sulfur can be reversed by a reductant, we added dithiothreitol (DTT) into the mixture of sulfane sulfur, AdpA, and the *P*_*wblA*_ probe (the DTT dosage was 2 folds of HS_n_H/S_8_). After DTT treatment, AdpA restored the high affinity with the *P*_*wblA*_ probe (Fig. 3), which had been attenuated by sulfane sulfur. These phenomena demonstrated that the affinity attenuation of AdpA to its cognate DNA caused by sulfane sulfur is reversible.

**Fig. 3.**
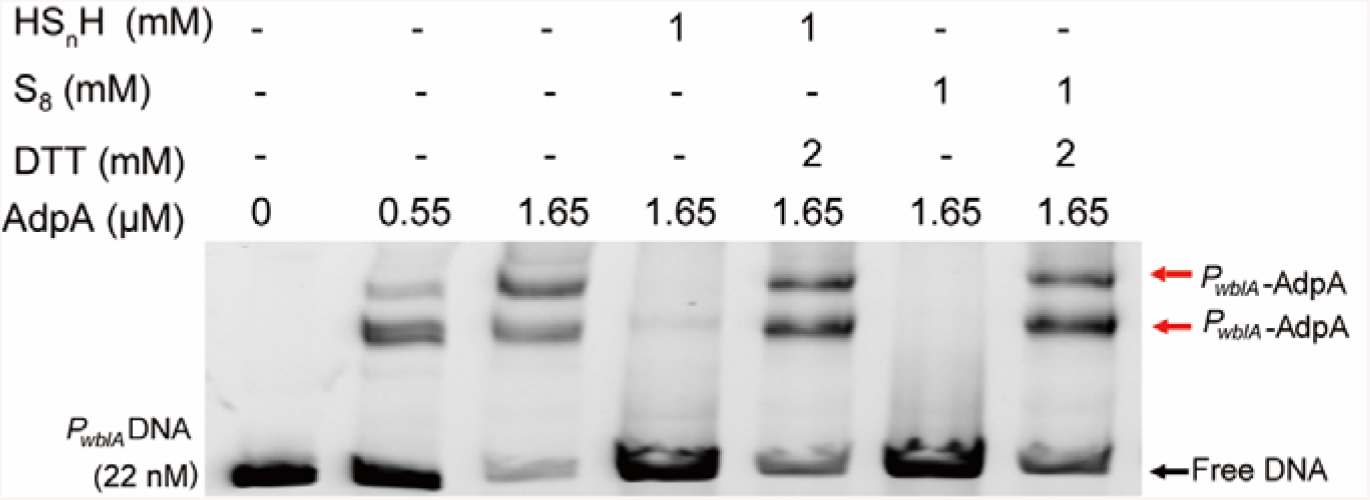
EMSA analysis of AdpA binding to *P*_*wblA*_ promoter DNA. All lanes contained 22 nM *P*_*wblA*_ DNA, lanes 2-7 contained AdpA, lanes 4 and 5 contained HS_n_H, lanes 6 and 7 contained S_8_, lanes 5 and 7 contained DTT.

### Sulfane sulfur also affects the transcription of AdpA itself

The *adpA* gene is transcriptionally self-controlled (28). There are five AdpA binding sites in the *P*_*adpA*_ promoter (Fig. 4A). We designed a pair of primers (*adpA-wt*) from the undeleted part of *adpA* gene (Fig. 4B) and used these primers to analyze the transcription change of *adpA* in wt and Δ*adpA*. The transcription level of undeleted part was ∼30-fold higher in Δ*adpA* than that in wt, indicating that in the absence of AdpA, the strength of *P*_*adpA*_ is higher, i.e., AdpA acts as a repressor for its own transcription (Fig. 4C)

**Fig. 4.**
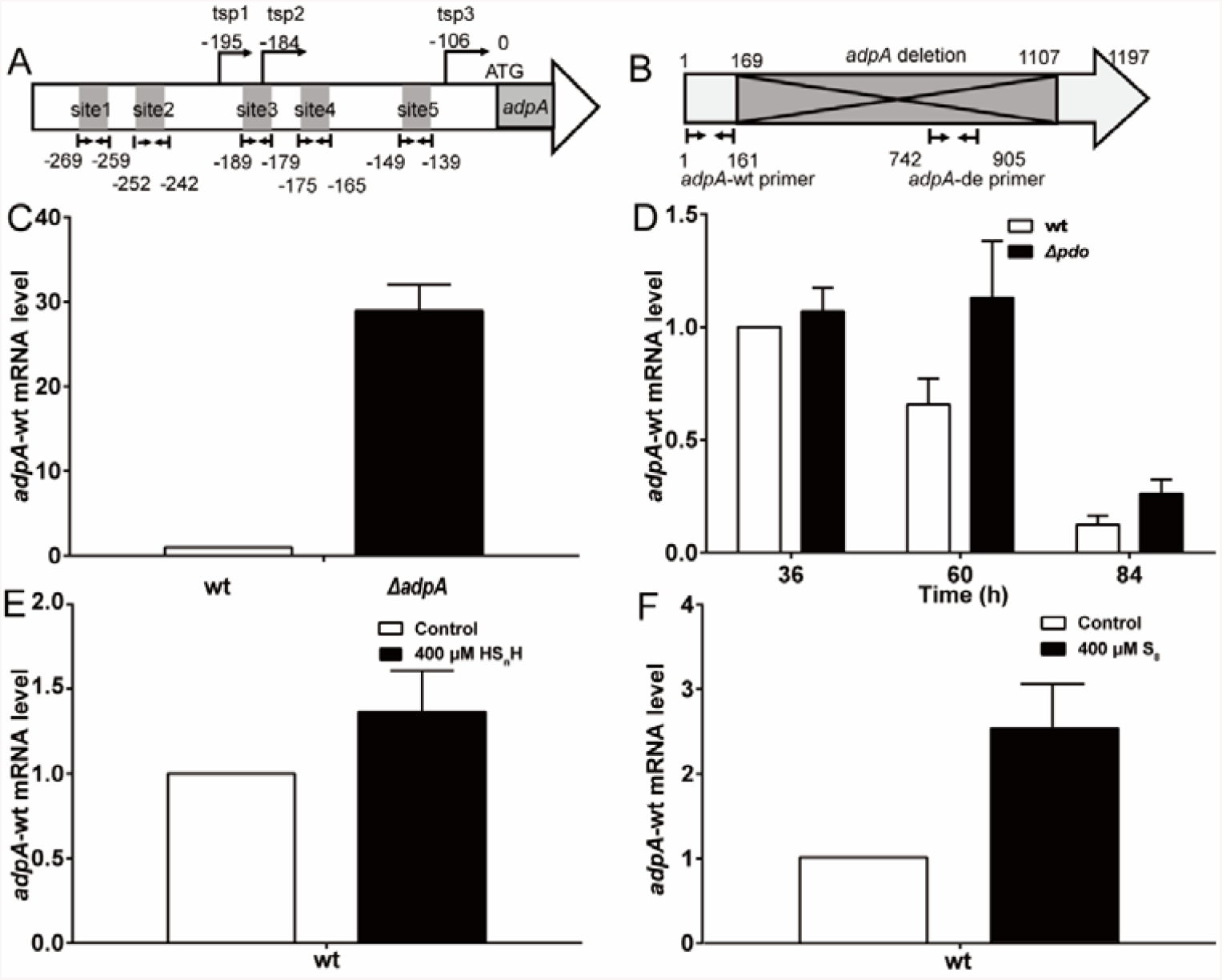
Sulfane sulfur affects the transcription of *adpA* itself. (A) Schematic diagram of the the AdpA binding sites in *adpA* promoter region. (B) Schematic diagram of the the AdpA coding sequence. The fragment covers 169 bp to 1107 bp was deleted in Δ*adpA*. The *adpA-*wt and *adpA-*de primers were used to test the undeleted and deleted sequences, respectively. (C) RT-qPCR analysis of *adpA-*wt mRNA level in wt and Δ*adpA*. (D) RT-qPCR analysis of *adpA-*wt mRNA level in wt and Δ*pdo*. The expression level in wt at 36 h was arbitrarily set to one. The y-axis shows the fold change of transcription levels in wt and Δ*pdo* over the levels in wt at 36 h. Data were from three independent repeats and shown as average ± SD. (E,F) RT-qPCR analysis of *adpA-*wt in wt and Δ*adpA* after inducing by HS_n_H (E) and S_8_ (F) for 1 h. Data were from three independent repeats and shown as average ± SD.

To test whether sulfane sulfur can affect this self-repression, we compared transcription levels of *adpA* in the wt stain and the Δ*pdo* strain. In the latter intracellular sulfane sulfur is accumulated due to lack of the persulfide oxidation gene (*pdo*) (23). Results showed that the *adpA* transcription levels were higher in Δ*pdo* than in wt (Fig. 4D). We then used exogenous sulfane sulfur to treat the wt strain and found that both HS_n_H and S_8_ can increase *adpA* transcription (Fig. 4E and 4F). EMSA showed that sulfane sulfur also reduced the affinity of AdpA to *P*_*adpA*_ probe, as the unbound probe increased after the addition of HS_n_H and S_8_ (Fig. 5A). Fluorescence polarization (FP) analysis was performed, and results showed that HS_n_H obviously increased the K_D_ value (the equilibrium dissociation constant) of AdpA to the *P*_*adpA*_ probe, as well as to the *P*_*wblA*_ probe (Fig. 5B and 5C), indicating that the affinity of AdpA to these promoters was attenuated by HS_n_H. Together, these results indicated that sulfane sulfur attenuates the self-repression of *adpA* transcription.

**Fig. 5.**
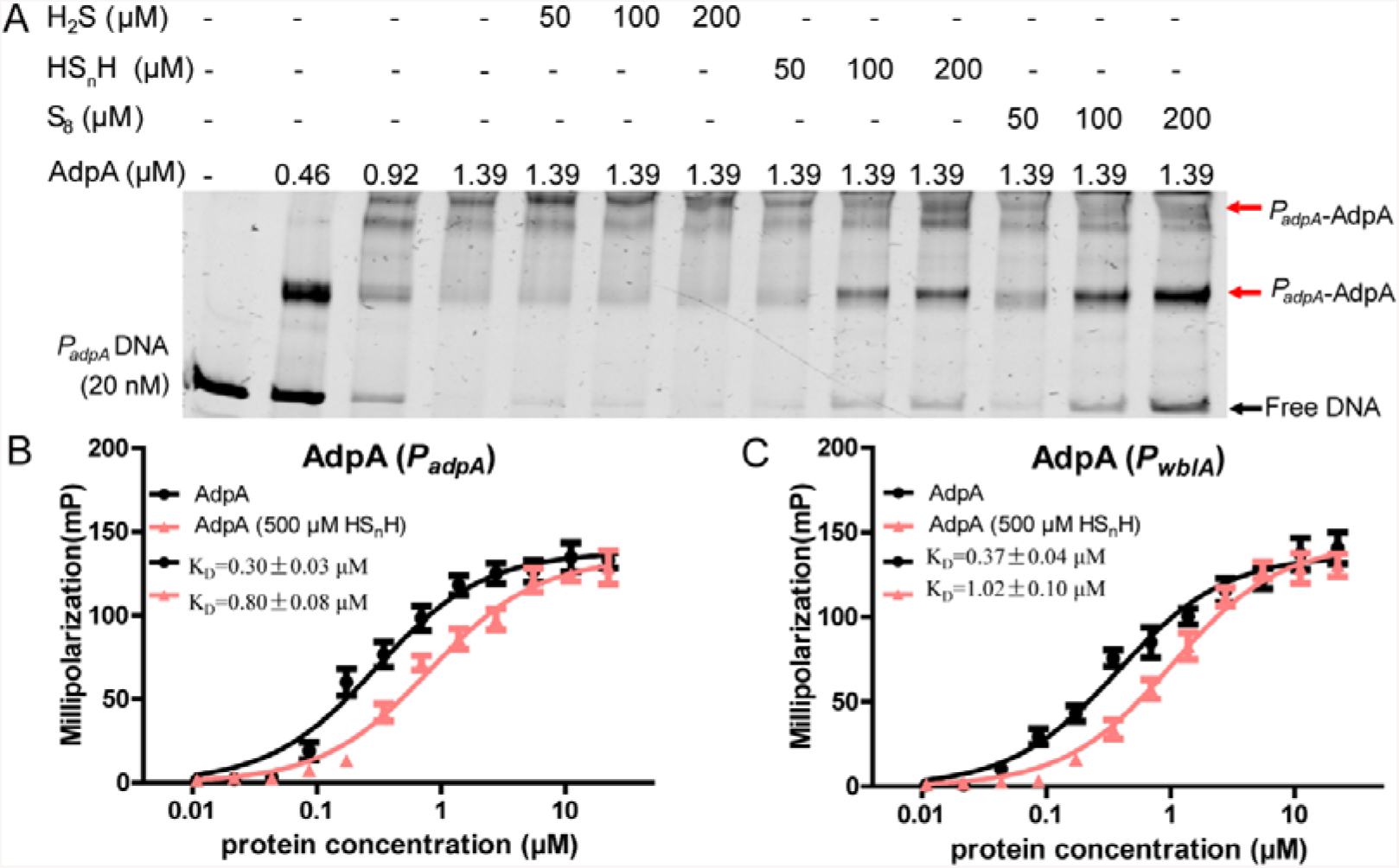
EMSA and FP analysis of AdpA binding to DNA probes. (A) EMSA analysis of AdpA binding to *P*_*adpA*_ probe. All lanes contained 20 nM probe, lanes 2-13 contained AdpA, lanes 5-7 contained H_2_S, lanes 8-10 contained HS_n_H, lanes 11-13 contained S_8_. (B,C) FP analysis of AdpA binding to *P*_*adpA*_ probe (B) and *P*_*wblA*_ probe (C). 1 nM FAM-labeled *P*_*adpA*_ or *P*_*wblA*_ was incubated with increasing amounts of AdpA or HS_n_H treated AdpA. The K_D_ values were calculated using GraphPad Prism 5 software. Data were from three independent experiments and shown as average ± SD.

### Kinetics simulation of the AdpA controlled promoter system

To better understand how sulfane sulfur influences the performances of *P*_*actII-4*_, *P*_*wblA*_, and *P*_*adpA*_, we used two mathematical models to simulate the dynamics of the promoter system. For simplification, we only took account of the influence of AdpA and sulfane sulfur on the system (Fig. 6). Equations and related parameters used for modelling are provided in supplementary materials. When cellular sulfane sulfur is low, *P*_*adpA*_ and AdpA compose a classic closed negative-feedback loop. Therefore, the *P*_*adpA*_ strength displays an oscillating dynamics, leading to oscillating AdpA expression. Since the expression of ActII-4 and WblA is activated by AdpA, their expression oscillates correspondingly with a short delay (Fig. 6A). When cellular sulfane sulfur is high, it attenuates the AdpA repression on *P*_*adpA*_ and breaks the negative-feedback loop. When the closed loop is totally broken, *P*_*adpA*_ is constantly on, leading to a constant level of AdpA and resulting in constant expression of ActII-4 and WblA (Fig. 6B). The variation of cellular sulfane sulfur between the low and high levels will affect the expression of *adpA, actII-4*, and *wblA*.

**Fig. 6.**
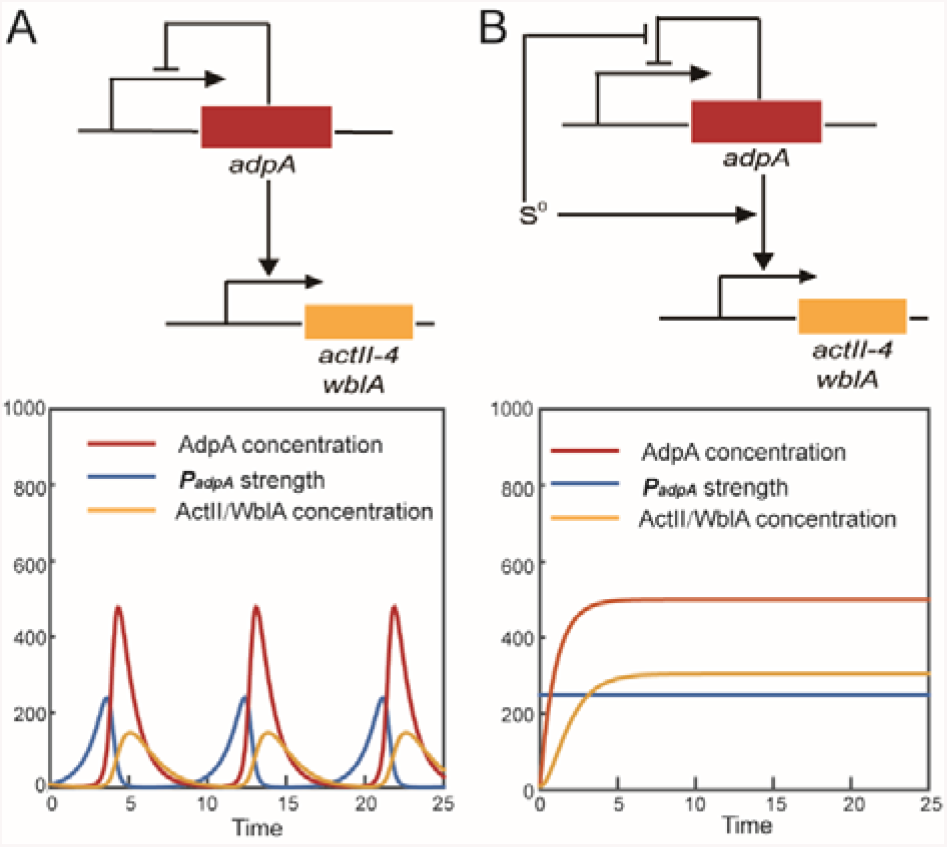
Modelling dynamics of the AdpA controlled promoters. (A) In the absence of sulfane sulfur, *P*_*adpA*_ is self-repressed by AdpA to form a negative-feedback loop, and hence both *P*_*adpA*_ strength and AdpA amount show an oscillating dynamics. (B) Sulfane sulfur breaks the negative-feedback loop, which leads to a constant expression of AdpA, ActII-4, and WblA. Equations and related parameters used for modeling are provided in supplementary materials.

### The cysteine residue Cys^62^ is critical for AdpA sensing sulfane sulphur

Sulfane sulfur can react with cysteine residues of certain proteins to change their configurations (29, 30). AdpA contains four cysteine residues, Cys^62^, Cys^126^, Cys^187^, and Cys^307^. To find out which cysteine residue involves in the AdpA-sufane sulfur interaction, we made cysteine-to-serine mutation on each cysteine residue of AdpA. The mutated *adpA* genes were complemented in the Δ*adpA* strain. When growing in YBP agar medium, the Δ*adpA*::*adpA*_*C126S*_, Δ*adpA*::*adpA*_*C187S*_, and Δ*adpA*::*adpA*_*C307S*_ strains did not show obvious difference from the wt and Δ*adpA*::*adpA* strains. However, the Δ*adpA*::*adpA*_*C62S*_ strain was distinct from others. It lost the ability of generating spores and its ACT production was also apparently lower (Fig. 7A). These phenotype changes indicated that Cys^62^ is critical for AdpA performing its regulatory function.

**Fig. 7.**
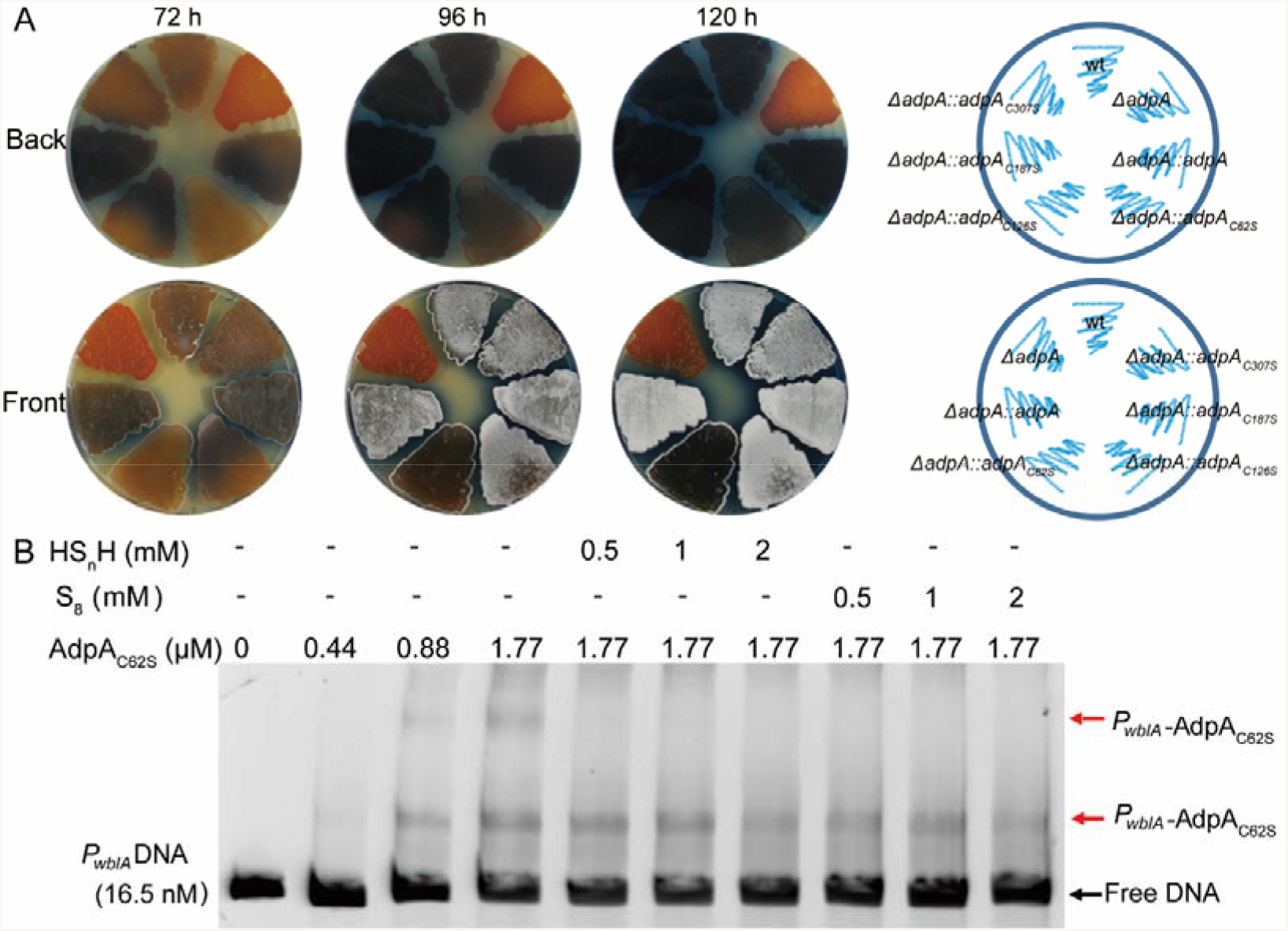
Cys^62^ residue is critical for AdpA sensing sulfane sulfur. (A) Phenotypes of wt, Δ*adpA* and complementary strains (Δ*adpA::adpA*_*C62S*_, Δ*adpA::adpA*_*C126S*_, Δ*adpA::adpA*_*C187S*_ and Δ*adpA::adpA*_*C307S*_) grown on YBP medium. Images were captured from both sides of the plates. (B) EMSA analysis of AdpA_C62S_ binding to *P*_*wblA*_ DNA. All lanes contained 16.5 nM probe DNA, lanes 2-10 contained AdpA, lanes 5-7 contained HS_n_H and lanes 8-10 contained S_8_. Black arrow indicates the free DNA probe, and red arrow indicates *P*_*wblA*_-AdpA_C62S_ complex.

EMSA was then performed to examine how C62S mutation affected the binding of AdpA to its cognate promoter. AdpA_C62S_ still bound to *P*_*wblA*_ DNA fragment, but the binding was no longer influenced by sulfane sulfur even when sulfane sulfur was added at high concentrations (>1000 fold higher than that of AdpA_C62S_) (Fig. 7B).

To check how sulfane sulfur reacts with AdpA, purified AdpA was treated with HS_n_H or DTT. The treated-AdpA was labeled with iodoacetamide (IAM) and then subjected to trypsin digestion, followed by LTQ-Orbitrap tandem mass spectrometry analysis. For the HS_n_H treated-AdpA, two peptides (1 and 2, Fig. 8) were identified. In peptide 1 (1299.67 Da), the Cys^62^ residue was directly blocked by IAM to form Cys^62^-AM (acetamide) (Fig. 8 and Fig. S1). In peptide 2 (1331.64 Da), a mass increase of 32 (+32 MW) was identified. MS^2^ spectrum indicated that the +32 MW happened on thiol group of Cys^62^ to form peptide-S-AM (Fig. 8 and Fig. S2). The MS^1^ signal intensity ratio of peptide 1/peptide 2 was 17 %, As the control, only a peptide with Cys^62^-AM (1299.67, peptide 3) was identified, corresponding to a direct blockage of IAM on Cys^62^ residue (Fig. 8 and Fig. S3). These results indicated that sulfane sulfur can modify Cys^62^-SH to form Cys^62^-SSH.

**Fig. 8.**
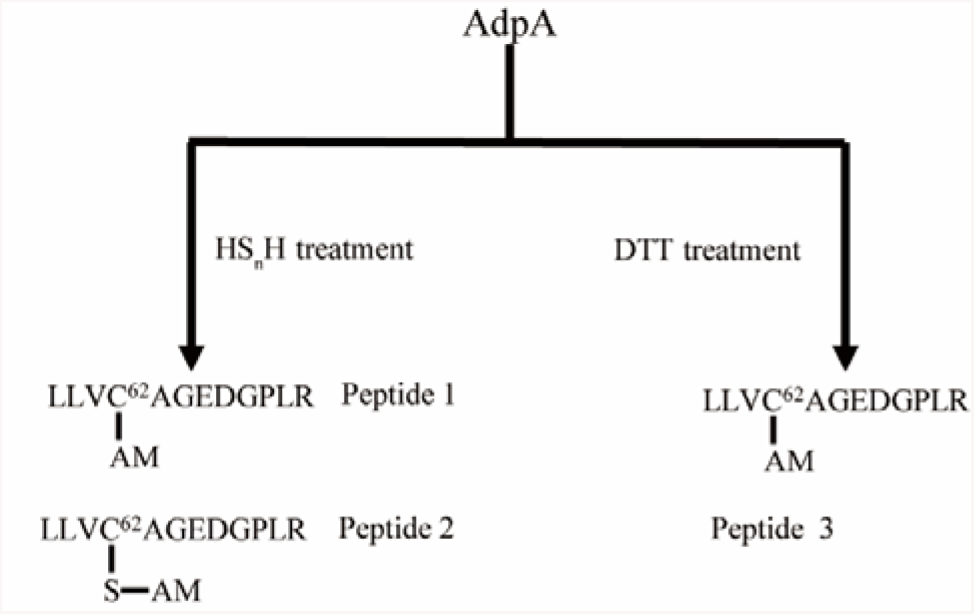
LC-MS/MS analysis of HS_n_H treated and DTT treated AdpA. MS^2^ data of the peptides are provided in Figs S1–S3.

### The thiol group of Cys^62^ is accessible to solution due to its location on the AdpA 3D structure

The 3D structure of AdpA was modelled by using AlphaFold 2 (https://cryonet.ai/af2/). The crystal structure of a truncated AdpAsg containing only the DNA binding domain, which is from *Streptomyces griseus*, is available in PDB database (PDB: 3w6v). We aligned AdpAsg with the modelled AdpA, and the alignment parameter RMSD was 0.365, indicating a high confidence of the predicted structure of AdpA (Fig. 9A and 9B).

**Fig. 9.**
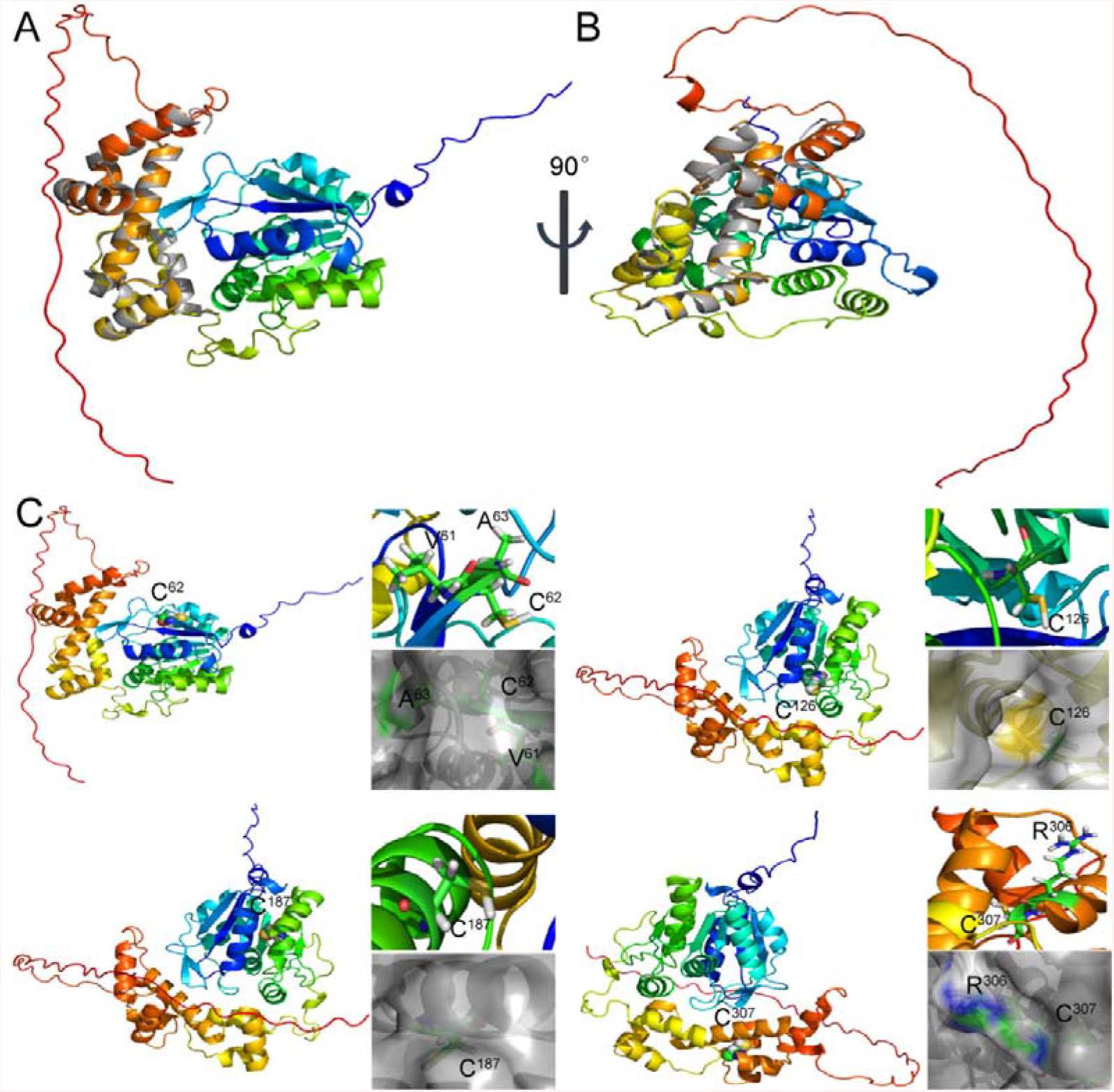
AlphaFold 2-predicted 3D structure of AdpA. (A, B) Alignment of the predicted AdpA structure (multicolor) with the AdpAsg crystal structure (grey). (C) Locations of the cysteine residues in AdpA. Yellow spheres represent the sulfur atoms.

We then analyzed locations of the four cysteine residues in AdpA (Fig. 9C). Cys^62^, Cys^126^, and Cys^187^ locate in the ThiJ/PfpI/DJ-1-like dimerization domain and Cys^307^ locates in the AraC/XylS-type DNA-binding domain (DBD). Cys^187^ and Cys^307^ fold into interior of AdpA, and hence they are protected from sulfane sulfur attack. Differently, Cys^62^ locates near the protein surface, and its thiol group is exposed to solution, which may explain why Cys^62^ can be modified by sulfane sulfur. It is noteworthy that Cys^126^ locates on the interior surface of a tunnel through the dimerization domain. Therefore, sulfane sulfur compounds may not enter this tunnel to react with its thiol group. In addition, the distance between any two cysteine residues is too far to form a disulfide (S-S) or tri-sulfide (S-S-S) bond.

### The cysteine residues are conserved in *Streptomyces* AdpAs

We analyzed AdpA and its homologues in *Streptomyces* genus. In the PATRIC database, 2752 of 3033 sequenced *Streptomyces* strains contain AdpA, accounting for a 90.73% prevalence. We selected some representative AdpA sequences to construct a phylogenetic tree. Results revealed that AdpA homologues were not on one evolutionary branch (Fig. S4). However, when we performed multiple sequence comparison with them, we found that their four cysteine residues were highly conserved, including Cys^62^ (Fig. S5). These results suggested that using cysteine residues to sense sulfane sulfur may be a common mechanism for AdpA functioning in *Streptomyces*.

## DISCUSSION

The global transcription factor AdpA plays an important role in regulation of secondary metabolism and morphological development in the *Streptomyces* genus (31-35). Its own expression is controlled by multiple factors. In this study, we discovered that sulfane sulfur affects AdpA activity via the post-translational modification in *S. coelicolor* M145. After reacting with sulfane sulfur, the affinity of AdpA to its cognate promoters, *P*_*adpA*_, *P*_*actII-4*_, and *P*_*wblA*_, is attenuated, which increases the expression of AdpA, *actII-4* and *wblA*. We constructed a simplified model to understand the effect of sufane sulfur on the AdpA-controlled promoters. As shown in our simulation (Fig. 6), *P*_*adpA*_ is under the control of a negative feedback loop of self-repression. Without presence of sulfane sulfur and/or other disturbing factors, activation of AdpA on *actII-4* and *wblA* expression cannot last long time due to the feedback. Sulfane sulfur modifies AdpA to break/remove the self-repression, and then the activation can last longer. Thus, the effect of sulfane sulfur on the AdpA regulon may represent a fine-tuned regulation for the production of antibiotics and morphological development. Further investigation is required to understand the fine regulations.

Furthermore, we found that Cys^62^ is critical for AdpA sensing sulfane sulfur. Sulfane sulfur treatment leads to a sulfhydration modification in Cys^62^ (Cys^62^-SSH). This modification is not observed in the other three cysteine residues of AdpA. The AlphaFold 2 predicted structure shows that the thiol of Cys^62^ is accessible to solution while the other thiols are not, which may explains why Cys^62^ is easily sulfhydrated by sulfane sulfur. However, it’s still unknown how this post-translational modification affects AdpA configuration. Since Cys^62^ locates in the dimerization domain, it may affect the dimerization and then alter the AdpA binding to its cognate DNA.

In recent years, a few transcription factors that can be modified by sulfane sulfur have been identified from different microorganisms. These transcription factors can be categorized into two groups. Group I consists of specific regulators for genes related to sulfur metabolism, including BigR (36), CstR (37), FisR (38), CsoR (23), and SqrR (39, 40). They control the expression of sulfane sulfur oxidation enzyme PDO and sulfane sulfur transferase RhoD (BigR is in this case). Since H_2_S oxidation enzyme SQR often locates in the same operon with PDO and RhoD, they also controls SQR expression (CstR, FisR, and SqrR are in this case) (41). Group I regulators can sense the intracellular level of sulfane sulfur via their cysteine residues: when sulfane sulfur level is high, cysteine residues are sulfhydrated to form RS_n_H (n≥2) or RS_n_R (n≥3) bond, which leads configuration change of the regulator and subsequently high expression of PDO and other genes, and then sulfane sulfur level is decreased through oxidizing to sulfite (19). Therefore, Group I regulators mainly function as managers to maintain the homeostasis of intracellular sulfane sulfur.

Group II includes global or multi-functional transcription factors currently including MgrA (21), MexR (42), and OxyR (43). MgrA is a global virulence regulator of *Staphylococcus aureus*. It sense intracellular level of sulfane sulfur to regulate the expression of virulence factors (21). MexR controls the multiple antibiotic resistance process in *Pseudomonas aeruginosa*, it senses intracellular sulfane sulfur to regulate the expression of *mexAB*-*oprM* multidrug efflux operon (42). OxyR is a global anti-oxidation regulator in many bacteria. Recently it was found that OxyR also senses sulfane sulfur and controls expression of sulfane sulfur reducing enzymes (43). Same as Group I, Group II regulators also sense sulfane sulfur via their cysteine residues.

AdpA is deemed as a Group II regulator since it senses sulfane sulfur and accordingly adjust the ACT production and spore formation in *S. coelicolor*. Bioinformatics analyses indicate that AdpA and its Cys residues are highly conserved in *Streptomyces* spp., and further investigations of this protein and its homologues should provide insights into how sulfane sulfur regulates the production of secondary metabolites and morphological developments in this genus. The widespread existence of AdpA implies that sulfane sulfur may play a wide range of regulatory functions in *Streptomyce*s, providing unlimited possibilities for sulfane sulfur as a signal molecule to stimulate increased production of important secondary metabolites, such as antibiotics, anti-tumor drugs, immunosuppressants, and antibiotics.

## MATERIALS AND METHODS

### Bacterial strains, plasmids and growth conditions

All strains and plasmids used or constructed in this work are summarized in Table S1.

*Streptomyces* strains were cultivated at 30°C on MS (mannitol soya flour) solid medium (44), YBP (yeast-beef-peptone) solid or liquid medium (45) were used for different experiments including spore suspension preparation, intergeneric conjugation, growth assay, RNA isolation and phenotypic observation. All *E. coli* strains were cultured at 37°C on solid or liquid Luria-Bertani (LB) medium. *E. coli* DH5α and *E. coli* BL21 (DE3) strains were used as hosts for plasmid construction and protein expression, respectively. *E. coli* ET12567 (pUZ8002) was used as a media for transferring nonmethylated DNA to *Streptomyces*. When required, ampicillin (100 μg/mL), apramycin (50 μg/mL), chloramphenicol (25 μg/mL), kanamycin (50 μg/mL), hygromycin (50 μg/mL), or nalidixic acid (25 μg/mL) were added into the medium.

### Preparation of sulfane sulfur species and other sulfur containing compounds

Sodium hydrosulfide (NaHS, H_2_S donor) and thiosulfate were purchased from Sigma Aldrich. S_8_ solution was freshly prepared by dissolving excess sulfur powder in acetone to reach saturation. The stock solution of HS_n_H (hydrogen polysulfides) was prepared by mixing sulfur powder, NaOH, and NaHS (40 mM of each chemical) in degassed distilled water at 30°C till the power was completely dissolved as previously described (42, 46). The concentrations of HS_n_H was determined with a cyanolysis method (47) and calibrated by using thiosulfate as the standard. The entire process was performed under anaerobic condition.

### Construction of *S. coelicolor* Δ*adpA*

All primers used in this experiment are listed in supplemental Table S2. The strain Δ*adpA* was constructed and generated using a homologous recombination method (48). Briefly, a 939 bp region was deleted from ORF of *adpA*, leaving the upstream168 bp (relative to the start codon) and the downstream 90 bp (relative to the stop codon) coding sequence of *adpA*. The knockout region was replaced by the *apramycin* resistance gene. The conjugation transfer was accomplished using the methylation-sensitive strain *E. coli* ET12567/pUZ8002 (containing the mutant plasmid pJTU-*adpA*) and *S. coelicolor* M145 following a previously reported protocol (49). The deletion mutant was verified by resistance screening and colony PCR with primers VeradpA-F/R.

### Construction of ΔadpA::adpA, ΔadpA::pMS82, ΔadpA::adpA_C62S_, ΔadpA::adpA_C126S_, ΔadpA::adpA_C187S_ and ΔadpA::adpA_C307S_

A DNA fragment carrying the *adpA* ORF (1197 bp) and its promoter (500 bp) was obtained using PCR amplification and connected to the ΦBT1 integrative vector pMS82 (50) to generate *adpA*-complemented plasmid pMS82-*adpA* (Table S1), which was then integrated into the *attP* site of Δ*adpA* genome by intergeneric conjugation. To construct negative control strains, empty pMS82 vector was also transformed into Δ*adpA*, these derivative strains were selected and confirmed by PCR and DNA sequencing.

In order to gain complementary strains with cysteine mutations, we adopted a point mutation strategy (51) based on the complementing plasmid pMS82*-adpA*, and obtained the resulting plasmids pMS82*-adpA*_*C62S*_, pMS82*-adpA*_*C126S*_, pMS82*-adpA*_*C187S*_ and pMS82*-adpA*_*C307S*_. The same method was used to obtain complementary strains Δ*adpA::adpA*_*C62S*_, Δ*adpA::adpA*_*C126S*_, Δ*adpA::adpA*_*C187S*_ and Δ*adpA::adpA*_*C307S*_. The primers used in this process are shown in Table S2.

### AdpA protein overexpression, purification and mutation

To construct the AdpA expression strain, the coding sequence of *adpA* was amplified from wt genomic DNA with primers ExadpA-F/R. The PCR product was purified and ligated into the pET15b vector with a C-terminal His tag to create plasmid pET-AdpA by using the ClonExpress™ II One Step Cloning Kit (TaKaRa). The plasmid was transformed into *E. coli* BL21 (DE3) cells, which were grown in LB medium at 37°C to an OD_600_ of about 0.6, and then a total of 0.5 mM isopropyl β-D-1-thiogalactopyranoside (IPTG) was added and an additional overnight cultivation was continued at 16°C. Cultures were collected by centrifugation and disrupted though a pressure cell homogenizer (SPCH-18) in sonication buffer (50 mM NaH_2_PO_4_, 250 mM NaCl, 20 mM imidazole, pH 8.0), 1 mM DTT was added before breaking the cells. Purification of the AdpA His-tagged proteins were performed with Ni-NTA-Sefinose column (Sangon) as described previously (23). The eluted fractions containing this protein were exchanged to a buffer (50 mM NaH_2_PO_4_, 50 mM NaCl, 10% of glycerol, pH 8.0) via PD-10 desalting column (GE), protein purification process was operated in an anaerobic glove box, the anaerobic box first exchanges gas with N_2_, and then the anaerobic box is filled with mixed gas (N_2_, 85%; H_2_, 10%; CO_2_, 5%) to keep the inside free of oxygen examined by a gas detector (ADKS-4, EDKORS, China).The purity of the protein was assessed by SDS-PAGE gel and its concentration was determined using the BCA Protein Assay Reagent (Thermo Fisher Scientific). AdpA cysteine mutant proteins (C62S, C126S, C187S and C307S) plasmids were constructed using the modified QuikChange™ method (51). We used the same method to express, purify and analyze these variants.

### Electrophoretic mobility shift assay (EMSA)

The DNA probes containing AdpA binding sequences were amplified from its target promoter region on the genomic DNA. Different sulfane sulfur compounds were reacted with the same amount of AdpA protein or cysteine mutant proteins in the binding buffer (20 mM Tris-HCI, 2 mM EDTA, 20 mM KCI, 0.5 mM DTT, pH 8.0) at room temperature for 20 minutes, and then the same amount of DNA was added to react at 30°C for 20 minutes, the samples were separated on an 8% nondenaturing polyacrylamide gel at 120V for 2 hour in ice (52), the gel was dyed with SYBR Green I (Sangon) for 20 minutes (38), All images were captured with a FlourChemQ system (Alpha Innotech).

### RNA preparation, RT-PCR and RT-qPCR

To extract RNA, spores (2×10^7^) of wt and Δ*adpA* strains were inoculated into the liquid YBP medium and incubated at 30°C with shaking (220 rpm) for 36 h to mid-exponential phase. HS_n_H (400 µM) or S_8_ (400 µM) was added. After 30 minutes of continued cultivation, these mycelia were harvested by centrifugation for 10 min, grounded into powder with liquid nitrogen. Similarity, the cultures of wt, Δ*adpA* and Δ*pdo* were collected on YBP plates at indicated times. All RNA were isolated with *SteadyPure* Universal RNA Extraction Kit (Accurate Biology) following manufacturer’s instructions, and their quality and concentration were determined by a NanoDrop ND-1000 (Thermo Fisher). Turbo DNaseI (Invitrogen) was utilized to remove genomic DNA. RT-PCR was carried out using a reverse transcriptase kit (Invitrogen) following manufacturer’s recommendations. RT-qPCR was performed using SYBR Premix Ex Taq (Takara). Roche LightCycler 480 thermal cycler was used to assess the melting curve and quantification of the PCR products, and detailed thermal cycling steps was carried out as the protocol previous described (53). The expression of *hrdB* mRNA was used as the internal standard to normalize the relative quantities of cDNA. The relative expression abundance of the target gene were analyzed using a relative quantification method of 2^-ΔCt (test gene-*hrdB*), three independent replicates were performed in parallel.

### Phenotypic analysis and actinorhodin production assay

*S. coelicolor* strains were cultured on solid YBP medium at 30°C for phenotypic observation and imaged at indicated times. ACT production measurement was performed as described previously (23, 54, 55). Briefly, *Streptomyces* strains were incubated on YBP medium for 7 or 10 days, mycelia were harvested from the plate. KOH (1 M final concentration) was added to treat the mycelia for 4 h. Then the mixtures were centrifuged to collect the supernatant, in which ACT concentration was determined by a spectrophotometer. Three independent biological experiments were replicated.

### Construction and testing of EGFP reporter systems

To construct the reporter plasmids, promoter fragments (nucleotides -400 to -1 upstream from gene translational start codon of *actII-4* and nucleotides -460 to -1 upstream from gene translational start codon of *wblA*) were amplified by PCR using primers pMS82-*actII-4*p-*egfp* S1-F/R and pMS82-*wblA*p-*egfp* S1-F/R (Table S2), respectively. Then these promoter fragments and a DNA fragment encompassing the *egfp* gene (coding for a green fluorescent protein from pEGFP-C2) were mixed and cloned into the pMS82 vector, yielding pMS82-*actII-4*p-*egfp* and pMS82-*wblA*p-*egfp*, respectively. Next, we introduced these reporter plasmids into wt and Δ*adpA* via conjugation with non-methylating *E. coli* ET12567 (pUZ8002), respectively. Exconjugants with hygromycin-resistant strains were screened and checked by PCR using specific primers (Table S2).

Strains containing the reporter plasmid were pre-cultured in liquid YBP medium for 36 h at 30°C. Subsequently, equal amount mycelia of each strain was transferred to the fluted bottle, and inducers (HS_n_H and S_8_) were added. After 60 min of induction, the bacteria were collected by centrifugation, and mycela were resuspended in 200 µl PBS buffer (OD_450_=2). EGFP fluorescence was measured using the microplate reader Synergy H1. The excitation wavelength and emission wavelength were set to 485 nm and 515 nm, respectively. The EGFP fluorescence intensity was normalized against cell density (fluorescence/OD_450_ of mycelia).

### LC-MS/MS analysis of AdpA

The analysis was performed following a previous report (23). Freshly purified protein AdpA was treated with 10 fold amounts of HS_n_H or DTT. After reacting at room temperature for 40 minutes. The reacted protein was treated with denaturing buffer (0.5 M Tris-HCl, 2.75 mM EDTA, 6 M guanadine-HCl, pH 8.0) containing 1 M iodoacetamide (IAM). The treatment was carried out in dark for 1 h, then sample was digested with trypsin (1:25, w/w) at 37°C for 20 h. The digestion products were filtered by C18 Zip-Tip (Millipore) and vacuum dried. Obtained peptides were resuspended in 10 μl ddH_2_O.

The Prominence nano-LC system (Shimadzu) equipped with a custom-made silica column (75 μm × 15 cm) packed with 3-μm Reprosil-Pur 120 C18-AQ was used. Positive electrospray ionization was performed and the ions were scanned via LTQ-Orbitrap Velos Pro CID mass spectrometer (Thermo Scientific); the data were analyzed by a data-dependent acquisition mode with Xcalibur 2.2.0 software (Thermo Scientific). Full-scan MS spectra (from 400 to 1800 m/z) was detected and assessed by the Orbitrap at a resolution of 60,000 at 400 m/z.

### AdpA structure modelling

The AlphaFold 2 algorithm (56) was used to predict the tertiary structure of AdpA. This method used the custom multiple sequence alignment (MSA) option and accessed by the Colab server on GitHub (https://github.com/sokrypton/ColabFold). Structural model of AdpA was analysed and visualized with PyMOL.

### Fluorescence polarization (FP) analysis

FP analysis experiments were performed following a reported protocol (57). DNA probes were amplified by PCR and labelled by 5’6-FAM (Carboxyfluorescein) (Sangon). Purified AdpA (treated with 0.5 mM HS_n_H for 10 min or not) was diluted to different concentrations (0.01 µM∼22.5 µM). The reaction buffer contained 10 mM Tris–HCl and 75 mM NaCl, pH 7.5. After mixing diluted AdpA and labelled DNA in the reaction buffer, the solution was incubated at 37°C for 15 min in the dark. The fluorescence was detected with BioTek Synergy HT. The K_D_ value was calculated using GraphPad Prism 5 software.

## ACKNOWLEDGEMENTS

This work was supported by the National Natural Science Foundation of China (91951202) and the National Key R&D Program of China (2018YFA0901200).

